# MLCK/MLCP regulates mammalian axon regeneration and redistributes the growth cone F-actin

**DOI:** 10.1101/2024.07.30.605892

**Authors:** Saijilafu, Wei-Hua Wang, Jin-Jin Ma, Yin Yin, Yan-Xia Ma

## Abstract

Axon regrowth is a key determinant of the restoration of the biological function of the nervous system after trauma. However, mature mammalian neurons have limited capacity for axon regeneration. We have previously demonstrated that neuronal axon growth both in the central and the peripheral nervous systems is markedly enhanced when non-muscle myosin II (NMII) is inhibited with blebbistatin. The activity of NMII is primarily regulated by MLCK and MLCP *via* the phosphorylation and dephosphorylation of its light chain, respectively; however, the functional roles of MLCK and MLCP in mammalian axonal regeneration remain unknown. In the present study, we provide strong evidence that the inhibition of MLCK activity significantly blocks axon regeneration in mice. Conversely, the inhibition of MLCP promotes axon regrowth of both the peripheral and central nervous system. Our findings further indicate that the MLCK/MLCP regulates axon regeneration and redistributes the growth cone F-actin, and this result suggests that direct regulation of the growth cone machinery is a potential strategy to promote axon regeneration.

## Introduction

The coordinated regulation of growth cone cytoskeleton components is essential for axon growth. Additionally, cytoskeletal proteins in the growth cone are converging targets of extracellular inhibitory molecules. It is now recognized that extracellular inhibitory molecules, such as myelin-based inhibitors and chondroitin sulfate proteoglycans (CSPGs) in glial scars are a major hurdle for successful axon regeneration. These extracellular inhibitory molecules typically trigger the activation of the RhoA/ROCK pathway, leading to phosphorylation and, consequently, the deactivation of cofilin, a protein that severs actin filaments and promotes their depolymerization. Cofilin deactivation results in the stabilization of the growth cone actin cytoskeleton protein, and, subsequently, growth cone collapses. Accordingly, in order to overcome those inhibitory substrates and promote axonal outgrowth, several recent studies have focused on manipulating growth cone cytoskeleton dynamics, and have obtained promising results^1–4^. Here, we hypothesized that the appropriate spatiotemporal regulation of the local cytoskeletal machinery in the growth cone can provide an alternative and effective approach for enhancing axon regeneration.

We have previously shown that the inhibition of non-muscle myosin II (NMII) activity can promotes axonal growth over inhibitory substrates, such as CSPGs and myelin, in both the central and peripheral nervous systems (CNS and PNS, respectively)^1^. Furthermore, this growth-promoting effect is achieved *via* the reorganization of microtubules (MTs) in the growth cone^1^. Myosin proteins constitute a superfamily of ATP-dependent motor proteins known primarily for their role in muscle contraction. In the nervous system, however, myosin II usually drives actin retrograde flow in the growth cone, while the actin cytoskeleton powers directional growth cone motility^5,6^. An increasing number of studies have shown that the actin cytoskeleton and its regulators also play a critical role in axon elongation and guidance. Myosin II is typically activated *via* the phosphorylation of the N-terminus of its light chain by myosin light chain kinase (MLCK) and deactivated through dephosphorylation by myosin light chain phosphatase (MLCP). An early study reported that the expression of MLCK, a calcium/calmodulin-dependent kinase, is increased during axonal outgrowth in goldfish retinal ganglion cells^7^. The overexpression of constitutively active MLCK in Drosophila CNS neurons increases myosin II activity and leads to impaired axonal outgrowth^8^. Zhang *et al.* demonstrated that MLCK inhibition attenuates 5-HT-dependent neurite outgrowth in bag cell neurons of the mollusk Aplysia^9^. Additionally, the activation of MLCP/MLC signaling in PC12 cells was also shown to promote neurite outgrowth^10^. Together, these reports suggested that the MLCK/MLCP regulates axon growth in non-mammalian organisms. However, to our knowledge, the functional roles of MLCK and MLCP during mammalian axon regeneration remain unknown.

In the present study, we found that sciatic nerve injury elevated MLCK protein expression, leading to increased levels of MLC phosphorylation in dorsal root ganglion (DRG) neurons. Furthermore, the pharmacological inhibition or genetic silencing of MLCK blocked axon regeneration in the PNS and CNS both *in vitro* and *in vivo*, whereas the pharmacological inhibition or transcriptional silencing of MLCP dramatically promotes axon regeneration. We further found that the MLCK/MLCP activity also affects F-actin redistribution in the growth cone. Taken together, our results indicated that MLCK and MLCP may regulate axon regeneration and redistribute the growth cone F-actin, and the direct manipulation of cytoskeleton proteins in the growth cone may be a promising strategy for enhancing mammalian axon regeneration.

## Materials and Methods

### Animals and reagents

Adult ICR (8–10 weeks old, 20–30 g) were used. BDA (D1956) and CTB labeled with Alexa Fluor 488 (CTB-488, C34775) were obtained from Life Technologies. ML-7 hydrochloride (110448-33-4) was purchased from Santa Cruz Biotechnology. PDBu (P1269) was acquired from Sigma-Aldrich. Blebbistatin (S7099) was obtained from Selleck Chemicals. Antibodies against βIII-tubulin (TUJ1, T8578), MLCK (SAB1300116), and GFAP (G3893) were purchased from Sigma-Aldrich. Antibodies against β-actin (#4970) and p-MLC (#3675) were acquired from Cell Signaling Technology. Actin-stain 555 phalloidin (Cat^#^ PHDH1) was purchased from Cytoskeleton, Inc. siRNAs designed to target mouse MLCK (siMLCK) (5′-GGCAAATACACCTGTGAAG-3′, 5′-TGGTCAAAGAAGGGCAGAT-3′, 5′-AGCCAAAGGGAGTCAACAT-3′, and 5′-TCACGACGGGAATGAGATT-3′) and mouse MYPT1 (siMYPT1) (5′-CTGTGGATATCTCGATATTGC-3′) were obtained from GenePharma (Shanghai, China). The scrambled siRNA was used as control.

### Cell culture

Adult DRG neurons were dissociated from 8 to 10-week-old ICR mice and cultured in Minimal Essential Medium (MEM) containing 5% (*v*/*v*) fetal bovine serum (FBS), 5-fluoro-2-deoxyuridine/uridine (20 μM), and penicillin/streptomycin. Embryonic cortical neurons (from E14.5 embryos) and hippocampal neurons (from E18.5 embryos) were cultured in neurobasal medium supplemented with penicillin/streptomycin, GlutaMAX, and B27 supplement. To investigate axonal growth on inhibitory substrates, glass coverslips were first coated with poly-D-lysine (100 μg/mL) for 2 h, and then coated again with myelin or 10 μg/mL CSPGs for 2 h, as previously described^1,11^.

For the cell replating experiment, DRG neurons were cultured for 3 days, resuspended in culture medium, replated on new coverslips, and finally re-cultured for a further 18 h to allow new axons to grow, as previously described^12,13^. Briefly, in treatments denoted as “before replating”, neurons were first grown in the presence of 10 μM ML-7 during the initial 3-day culture period, and, after washing out the ML-7, they were replated and grown in the absence of drug(s) for 18 h. Treatments denoted as “after replating” refer to procedures in which neurons were first grown in the absence of a drug or drugs during the initial 3-day culture period, after which 10 μM ML-7 was added immediately after replating, followed by 18 h of culture in the presence of drug(s).

### Sciatic nerve axotomy

The sciatic nerve was transected at the sciatic notch using ophthalmic scissors. After surgery, the wound was closed, and the mice were allowed to recover. The mice were then euthanized and L4–L5 DRGs were isolated for cell culture, quantitative reverse transcription-PCR (qRT-PCR), and western blotting.

### siRNA transfection

Cells were transfected with siRNA by electroporation in accordance with the manufacturer’s instructions. Briefly, dissociated neurons were suspended in 80-µL solutions of Amaxa electroporation buffer containing siRNAs (2–3 nmol) and/or an EGFP-expressing plasmid (5 mg). The cells were then transferred to a 2-mm cuvette and electroporated using an Amaxa Nucleofector apparatus. After electroporation, the cells were immediately transferred to 500 µL of prewarmed culture medium and cultured on glass coverslips coated with 100 mg/mL poly-D-lysine. After 4 h, when the neurons were fully attached to the coverslips, the remaining electroporation buffer was discarded, and fresh culture medium (500 mL) was added. We used the pCMV– EGFP–N3 as control, and the pCMV– EGFP–N3 plasmid was from Clontech, Inc.

### qRT-PCR and western blotting

Total RNA was isolated using Trizol Reagent and assessed for integrity and concentration in a NanoDrop 1000 spectrophotometer. Each sample was then reverse transcribed using Maxima H Minus Reverse Transcriptase. Real-time qPCR was performed using SYBR Premix ExTaq II in a CFX96 Real-Time qPCR Detection System (Bio-Rad). Glyceraldehyde 3-phosphate dehydrogenase (*Gapdh*) was used as internal control. The sequences of the forward and reverse primers were MLCK: 5′-AGAAGTCAAGGAGGTAAAGAATGATGT-3′ and 5′-CGGGTCGCTTTTCATTGC-3′, respectively; and GAPDH: 5′-AGGTCGGTGTGAACGGATTTG-3′ and 5′-TGTAGACCATGTAGTTGAGGTCA-3′, respectively.

For western blotting, samples were homogenized in RIPA buffer. After boiling for 5 min, equal amounts of extracted protein (20 µg/sample) were separated using 10% sodium dodecyl sulfate–polyacrylamide gel electrophoresis, and electro-transferred to polyvinylidene fluoride membranes (Immobilon-P; Millipore). After blocking with 0.01 M PBS containing 5% (*w*/*v*) skimmed milk overnight, the membranes were incubated first with primary antibodies and then with the appropriate secondary peroxidase-conjugated antibodies. Protein bands were developed using ECL Prime Western Blotting Detection Reagent and subsequently quantified using Image Lab software. An anti-β-actin antibody was used as a loading control.

### Immunohistostaining

Mice were deeply anesthetized and transcardiacally perfused with 4% paraformaldehyde (PFA). L4–L5 DRG tissues were isolated, further fixed in 4% PFA, cryosectioned into 12-μm slices, and blocked with 5% FBS containing 0.3% Triton X-100. The sections were then incubated with the indicated primary antibody (monoclonal anti-βIII-tubulin antibody and/or polyclonal anti-MLCK antibody). Immunoreactivity was visualized using the appropriate fluorescently labeled secondary antibody. Alternatively, cultured DRG neurons were fixed in 4% PFA, blocked with 2% BSA and 0.1% Triton X-100, and then stained as described above. To visualize F-actin in the growth cones, *in vitro*-cultured neurons were stained with actin-stain 555 phalloidin following the manufacturer’s instructions.

### Axonal growth and growth cone quantification

All images were acquired at 1388 × 1040-pixel resolution using a Zeiss fluorescence microscope with AxioVision 4.7 software (Carl Zeiss Micro Imaging, Inc.) equipped with a 10×or a 20×objective. To quantify axon length, neurons with axonal processes exceedingly twice the diameter of their cell body were selected, and the longest axon from 100 neurons for each condition was measured using the curve tool in AxioVision 4.7. Meanwhile, growth cone size and F-actin area were measured using the outline tool in AxioVision 4.7. The size of the growth cone was ascertained from the neck of the microtubule to the edge of the axon tip and the F-actin area was determined from the hillock to the most distant edge of the growth cone.

### *In vivo* electroporation of DRGs and sciatic nerve crush

*In vivo* electroporation of adult mouse DRGs was performed as previously described^14^. Briefly, each mouse was anesthetized *via* an intraperitoneal injection of a ketamine (100 mg/kg) and xylazine (10 mg/kg) solution. The transverse process of L4 and L5 was carefully removed to expose the DRG tissue. A 1 μl solution containing indicated siRNAs together with the plasmid encoding EGFP (pCMV–EGFP–N3) was then microinjected into the L4–L5 DRG tissue using a Picospritzer II Microcellular Injection Unit followed by electroporation using a BTX ECM830 Electro Square Porator (five 15-ms pulses, 35 V, 900-ms interval). Following this, the muscle and skin were carefully sutured using 5-0 nylon sutures. The mouse was placed on a heated blanket (35°C) until it had completely recovered from anesthesia. Two days after electroporation, the sciatic nerves were crushed with forceps #5 and the crush sites were marked with an 11-0 nylon epineuria suture. After 3 more days, the mice were euthanized by perfusion of 4% PFA, and the whole sciatic nerves were dissected out and postfixed overnight in 4% PFA at 4°C. Prior to the whole-mount flattening process, it was essential to verify that the location of the epineural suture corresponded with the site of injury. Only experiments where the crush site was distinctly recognizable were considered for inclusion in the subsequent analysis. To quantify axon regeneration, all discernible axons labeled with EGFP within the sciatic nerve were manually traced and measured from the crush site to the distal tip.

### Optic nerve injury

The optic nerves of adult C57BL/6 mice were carefully exposed and crushed with a jeweler’s forceps #5 just behind the eyeball for 1 s. A small piece of gelatin sponge fully soaked with PBS or 50 μM PDBu was placed at the lesion site; alternatively, 2 μL of PDBu (50 μM) was intravitreally injected into the vitreous body. Simultaneously, 2 μL of CTB-488 was injected into the vitreous body of each animal after the optic nerve had been crushed. All microinjections were performed using the Picospritzer II Microcellular Injection Unit (Paker Ins.; pressure: 15 psi, duration: 6 ms). After 3 days, the optic nerves were dissected out, fixed in 4% PFA overnight at 4°C, and cut into a series of 12-μm longitudinal sections. The numbers of CTB-labeled axon fibers were counted at distances of 50 and 100 μm from the injury site. The length of the longest CTB-labeled axon fiber of each nerve was also measured.

### Spinal cord crush injury and PDBu injection

The T8 spinal cord was exposed under a microscope, the whole spinal cord was crushed for 2 s with a modified jeweler’s forceps #5 as previously described^15^, and the muscle and skin were separately sutured. A 100-µL volume of PDBu (50 μM) or DMSO was administered every 4 days by intrathecal injection. After 14 days, to trace and visualize the corticospinal tract axon, 1.6 μL of 10% BDA was injected into 4 sites of the sensorimotor cortex (1.0 mm lateral; 0.5 mm deep into the cortex; 1.0, 0.5, −0.5, and −1.0 posterior to bregma) with a 5 μL Hamilton syringe^16^. Fourteen days after BDA injection, mice were transcardically perfused with 4% PFA. The whole spinal cord was isolated from each mouse and post-fixed in 4% PFA for 6 h at 4°C. After dehydration using an increasing gradient of sucrose concentrations, the spinal cord was cut into 25-μm-thick sagittal sections, which were then stained with Cy3-streptavidin for 3 h following the manufacturer’s protocol. The lengths of regenerating axons were determined by measuring the distance from the lesion site to the tip of the BDA-labeled axon tip. The final axon length was the average calculated from 5 sections for each spinal cord. To quantify the retraction bulb, both maximum width of the enlarged distal tip of the axon and the width of its immediately adjacent axon shaft were measured. Then, the ratio of these two widths was then calculated. An axonal tip was considered as a retraction bulb if its tip/shaft ratio exceeded 4. The average number of retraction bulbs were calculated from 3 sections in every mouse for each group. Five mice were used for each condition.

### BBB locomotor scale score assessments

BBB score assessments were performed as previously described^17^. Briefly, mice were individually placed in an open, quiet space (45 ×30 cm) for 5 min while two individuals blinded to the experiments scored the hindlimb motor function of each animal. Assessments were made on day −1 (the day before injury) and on days 1, 3, 7, 14, 21, 28, 35, and 42 post-injuries for both the DMSO and PDBu treatment groups.

## Results

### The MLCK expression increased in adult sensory neurons following peripheral axotomy

Following injury, the expression of axon regeneration-related genes is usually upregulated. Thus, we first measured MLCK protein levels in DRG neurons following sciatic nerve transection. Compared with naive neurons, MLCK protein expression was significantly increased in DRG neurons 3 days after sciatic nerve transection (Figure 1A, B). The mRNA levels of MLCK in DRG neurons were also increased on days 1, 3, and 7 after axotomy, with the highest expression being observed on day 3 (Figure 1C). Given that MLCK phosphorylates MLC, we subsequently assessed the level of MLC phosphorylation and found that it was higher in DRG neurons following nerve injury than in control DRG neurons, likely due to the increase in MLCK expression (Figure 1A, D). Immunohistochemical staining further confirmed that both MLCK expression and the levels of phosphorylated MLC (p-MLC) increased in DRG neurons following sciatic nerve axotomy (Figure 1E, F). Taken together, these results indicated that MLCK is activated in DRG sensory neurons following peripheral axotomy, which leads to an increase in MLC phosphorylation.

**Figure 1.**
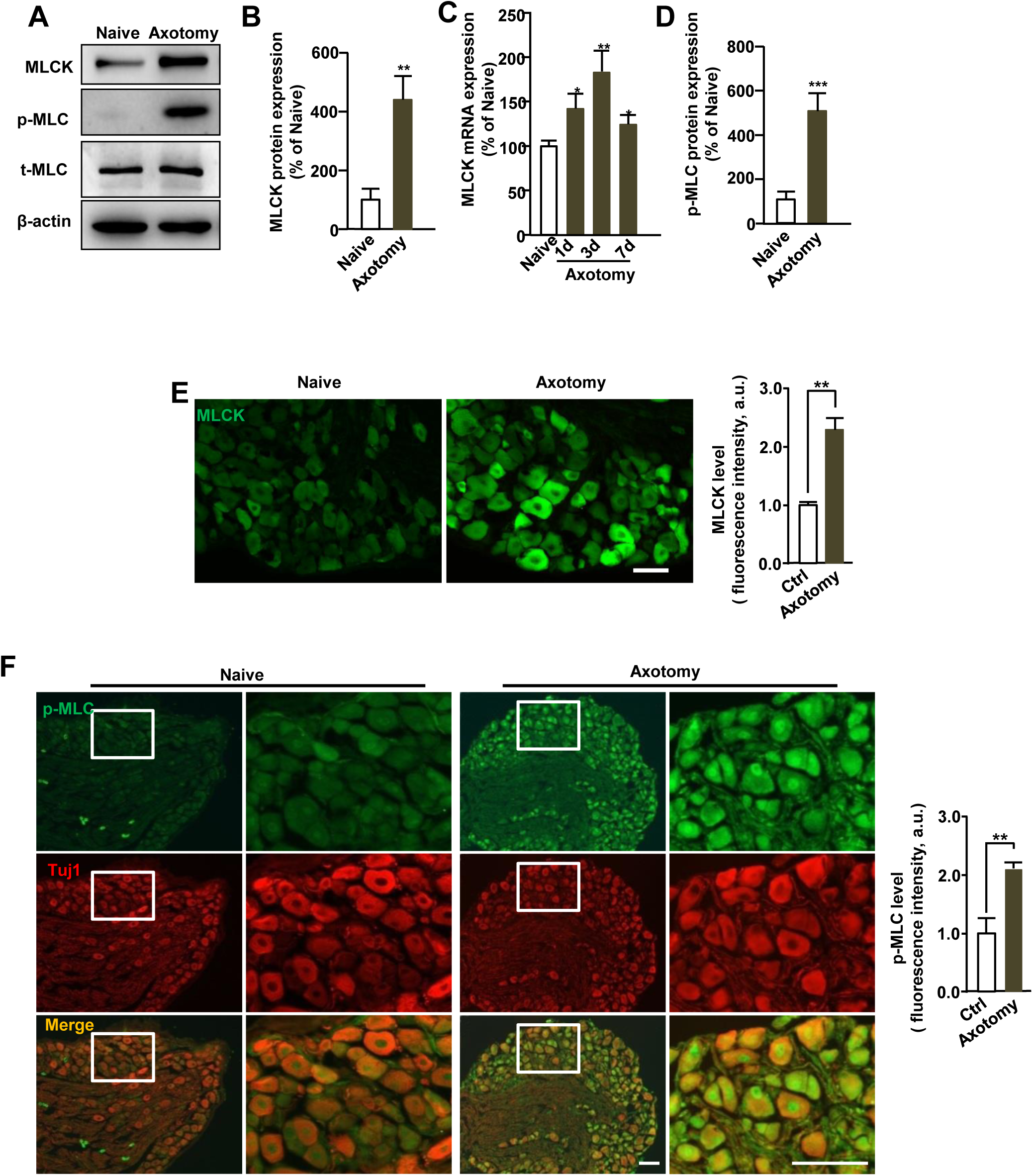
MLCK and phosphor-MLC expression increased in sensory neurons after peripheral axotomy. (A-D) Sciatic nerve axotomy was performed 3 days before and L4-L5 DRG tissues were collected for Western blotting or qPCR to examine the expression level of MLCK and phosphor-MLC (p-MLC). Representative images (A) and quantification of Western blots (B, D) demonstrate that expression levels of MLCK and p-MLC increase significantly in adult DRGs following sciatic nerve transection. (N=3, ** *p* < 0.01; *** *p* < 0.001). (C) L4-L5 DRGs were collected for qRT-PCR, 1, 3 and 7 days after axotomy, respectively. The results reveal that mRNA expression of MLCK in the DRG neurons also increased after axotomy after 1, 3 and 7 days, although the difference at 3 days was the most significant. (N=3, \**p* < 0.05; ** *p* < 0.01). (E-F) Sections of L4-L5 DRGs from naive and sciatic nerve axotomized mice were stained with antibodies against MLCK, Tuj-1 or p-MLC. Immunohistochemistry images indicate that MLCK (E) and p-MLC (F) expression were elevated in the sensory neurons following sciatic nerve axotomy. Scale bar: 100μm.

### The inhibition of MLCK impaired axon regeneration in peripheral sensory DRG neurons

Next, to explore the functional role of MLCK during peripheral axon regeneration, ML-7 (10 μM), a specific pharmacological inhibitor of MLCK, was added to DRG neuron cultures for 3 days. The results showed that ML-7 administration led to a significant reduction in MLC phosphorylation levels (Figure 2A, B, C) and impaired axonal growth in sensory neurons (Figure 2D, E), and these effects were dose-dependent (Supplementary Figure S1). The proportion of neurons with axons also decreased after ML-7 treatment (Figure 2E). Furthermore, the transfection of a small interfering RNA (siRNA) targeting MLCK (siMLCK) into dissociated DRG neurons significantly inhibited MLCK expression and reduced p-MLC levels (Figure 2F, G), while concomitantly blocking axon growth in adult DRG sensory neurons (Figure 2H, I). Next, using our previously described *in vivo* DRG electroporation technique^14^, we further explored the functional role of MLCK in sciatic nerve axon regeneration *in vivo*. We co-transfected siMLCK and an EGFP-expressing plasmid into lumbar vertebrae 4 and 5 (L4 and L5) DRGs of adult mice by electroporation. After 2 days, mice were subjected to sciatic nerve crush injury, and, after another 3 days, the lengths of regenerating axons were measured in whole-mount sciatic nerves. Our data showed that the mean length of *vivo* regenerating axons was significantly reduced in siMLCK-treated mice compared with that in control animals (Figure 2J, K). These findings suggested that MLCK activity is required for mammalian peripheral axon regeneration both *in vitro* and *in vivo*.

**Figure 2.**
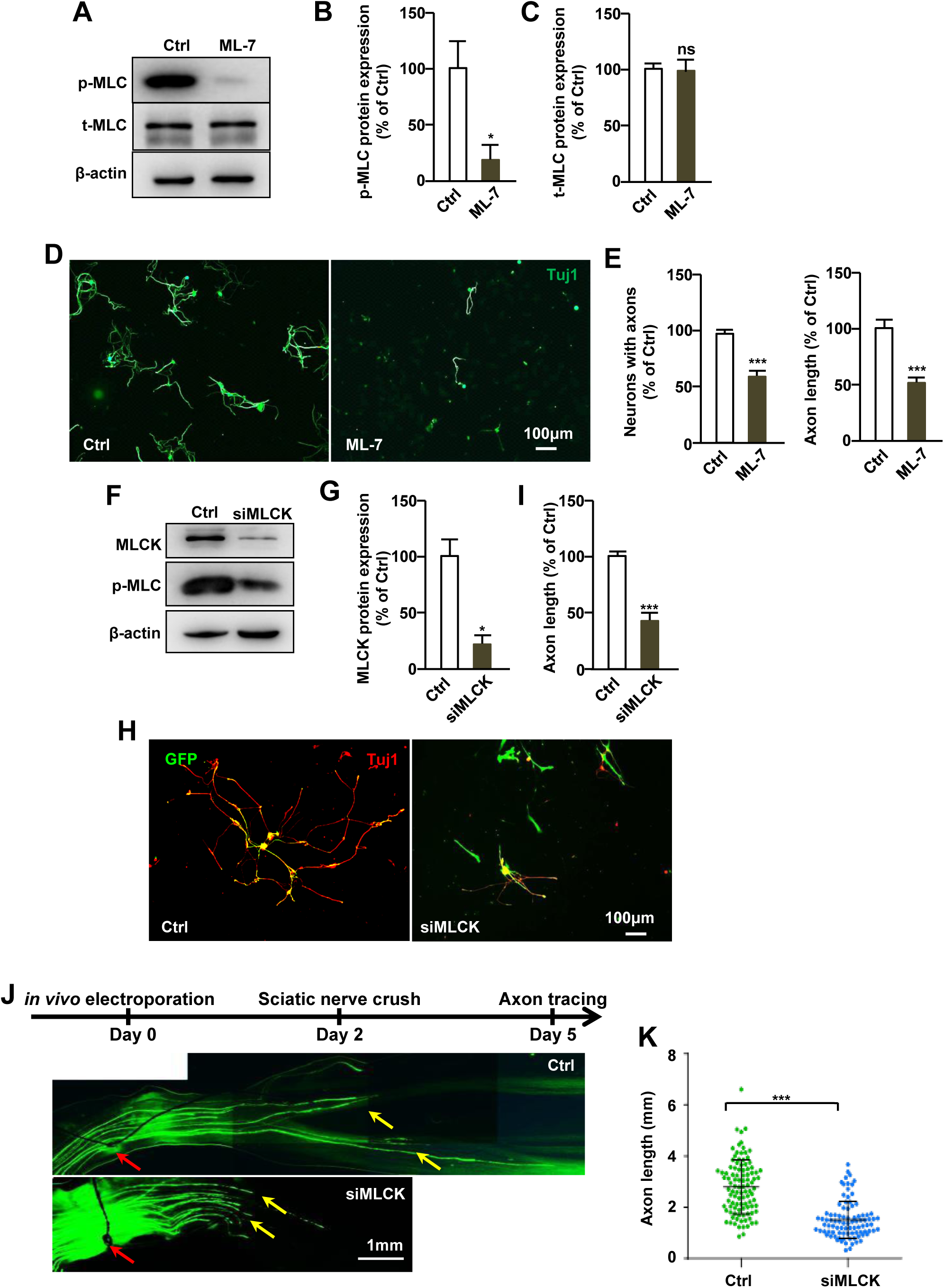
Suppression of MLCK activity inhibits axon growth in adult DRG neurons. (A-E) DRG neurons from adult mice were cultured and treated with DMSO or 10.0 µM ML-7. After 3 days, neurons were lysed for Western blotting or staining with Tuj1 antibody. Western blot images (A) and quantification of p-MLC (B) indicate that ML-7 treatment successfully inhibits MLCK activity. Quantification of axon length indicated that inhibition of MLCK activity with ML-7 dramatically blocks adult sensory neuronal axon growth *in vitro* (D, E), and reduces the numbers of neurons with axons (E). Scale bar: 100μm (N=3, \**p* < 0.05; *** *p* < 0.001). (F-I) Adult DRG neurons were isolated and electroporated with scramble siRNA (Ctrl) or MLCK siRNA (siMLCK) mixed with GFP. Tuj1 (Red) antibody was used to stain neurons 3 days after transfection. A representative image (F) and quantification of Western blots for MLCK (G) indicate that siMLCK significantly suppressed MLCK and p-MLC expression (N=3, \**p* < 0.05). Quantification of mean axon length (H, I) indicates that transfection of siMLCK inhibits adult DRG neuronal axon growth. Scale bar: 100μm. (N=3, *** *p* < 0.001). (J, K) L4/L5 DRGs of adult mice were electroporated with scramble siRNA (Ctrl) or siMLCK combined with GFP, respectively. Two days later, sciatic nerves were crushed and then regenerated axons traced after a further three days. Representative images (J) and measurement of regenerating axonal length (K) indicated that decreased MLCK expression with siRNA inhibits sensory axon regeneration *in vivo*. Crush sites are marked using red arrows, the marking suture was placed immediately after the crush injury. Regenerating axon tips are marked with yellow arrow (N=5, * *p* < 0.05). Scale bar: 1mm.

### The inhibition of MLCP promoted peripheral axon growth over inhibitory substrates

It is well known that the opposite roles of MLCK and MLCP in regulating the MLC phosphorylation status. Consequently, we also investigated the functional role of MLCP during axon regeneration. It has been reported that phorbol 12,13-dibutyrate (PDBu) can block MLCP activity, which leads to an increase in MLC phosphorylation^18^. Here, the results demonstrated that PDBu treatment (20 µM) significantly increased the p-MLC level in cultured DRG neurons (Supplementary Figure S2A, B). Importantly, we found that this occurred in parallel with greatly enhanced axonal growth in DRG neurons cultured on permissive substrates, such as poly-D-lysine (Supplementary Figure S2C, D). A key reason why CNS neurons cannot spontaneously regenerate axons after injury is due to the inhibitory local microenvironment. Myelin and CSPGs are well-characterized CNS-based inhibitory substrates that block axon regeneration. Therefore, we next examined whether inhibition of MLCP activity with PDBu promoted axon regeneration in the presence of these potential CNS-based inhibitory substrates. When dissociated adult DRG neurons were cultured on myelin extracts or CSPGs, sensory neuronal axon growth was significantly suppressed (Supplementary Figure S2C, D). Interestingly, however, when PDBu was added to the culture medium, axon growth was significantly enhanced on either substrate (Supplementary Figure S2C, D). This result indicated that PDBu treatment relieved the inhibitory effect of myelin and CSPGs on axonal growth in DRG neurons. MLCP is a phosphatase complex comprising PP1δ, MYPT1, and M20^19^. In this study, a siRNA specifically targeting MYPT1 (siMYPY1) was found to markedly increase MLC phosphorylation in cultured DRG neurons (Figure 3A, B). In line with the results obtained with the pharmacological inhibitor, siMYPT1 also significantly stimulated axonal growth in DRG sensory neurons cultured on myelin or CSPGs (Figure 3C, D). Furthermore, the transfection of siMYPT1 into L4–L5 DRGs of adult mice by electroporation also significantly promoted sciatic nerve axon regeneration *in vivo* (Figure 3E, F). Combined, these data indicated that the inhibition of MLCP activity enables mammalian axon regeneration in vitro and in vivo.

**Figure 3.**
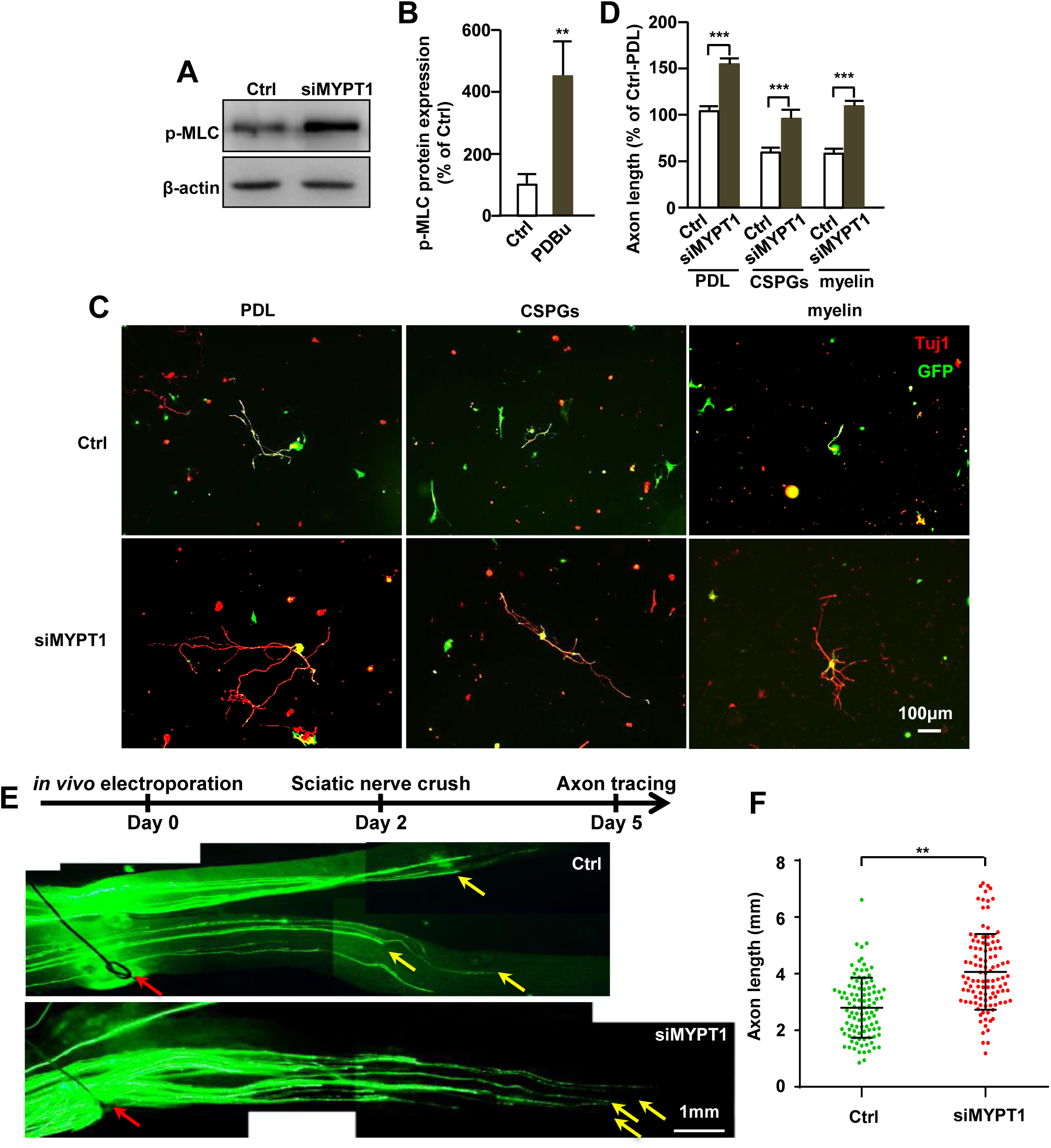
Knockdown of MLCP protein promotes axon growth from adult peripheral sensory neurons. (A-D) Adult DRG neurons were isolated and electroporated with scrambled siRNA (Ctrl) or MYPT1 siRNA (siMYPT1) mixed with GFP. A representative Western blot image (A) and quantification of protein band (B) indicate that transfection of siMYPT1 in cultured DRG neurons significantly increases p-MLC expression (N=3, ** *p* < 0.01). Quantification of axon length (C, D) indicated siMYPT1 transfection in the adult sensory neurons promotes its axon growth on the permissive substrate PDL or inhibitory substrates, such as CSPGs and myelin. (N=3, \*\*\**p* < 0.001). (E, F) L4/L5 DRGs of adult mice were electroporated with scrambled siRNA (Ctrl) or siMYPT1, respectively. Two days later, sciatic nerves were crushed and then regenerated axons traced after a further three days. Representative images (E) and measurement of regenerating axonal length (F) indicated that siMYPT1 promotes sensory axon regeneration *in vivo*. Crush sites are marked using red arrows, and the marking suture was placed immediately after the crush injury. Regenerating axon tips are marked with yellow arrow (N=5, * *p* < 0.05). Scale bar: 1mm.

### MLCK/MLCP activity regulated axon growth in the embryonic CNS in vitro

Compared with neurons of the PNS, those of the CNS have intrinsically poor axonal growth capability, which also helps explain why CNS neurons cannot regenerate axons spontaneously after injury. Therefore, we next examined the role of MLCK/MLCP on CNS axon regeneration using in vitro cultured cortical and hippocampal neurons from embryonic day (E)14.5 and E18.5 mice, respectively. To block MLCK activity, the neurons were treated with 10 µM ML-7 for 3 days. The results showed that the MLC phosphorylation levels were markedly inhibited with ML-7 treatment (Figure 4A); simultaneously, the mean axon length of ML-7-treated E14.5 cortical neurons was significantly reduced compared with that of the control neurons (Figure 4C, D). Furthermore, consistent with the pharmacological results, the siRNA-mediated suppression of MLCK expression in E14.5 cortical neurons (Figure 4B) significantly inhibited their axonal growth (Figure 4E, F). Conversely, the blockade of MLCP activity with PDBu (20 µM; Figure 4A) notably augmented cortical neuronal axonal growth (Figure 4C, D). The siRNA-mediated transcriptional silencing of MYPT1 expression also upregulated p-MLC levels in cortical neurons (Figure 4B), and, accordingly, also enhanced axonal growth (Figure 4E, F). Similar results were obtained with E18.5 hippocampal neurons (Figure 4G-J). Overall, these findings demonstrated that MLCK/MLCP activity also regulates axon growth in the developing CNS.

**Figure 4.**
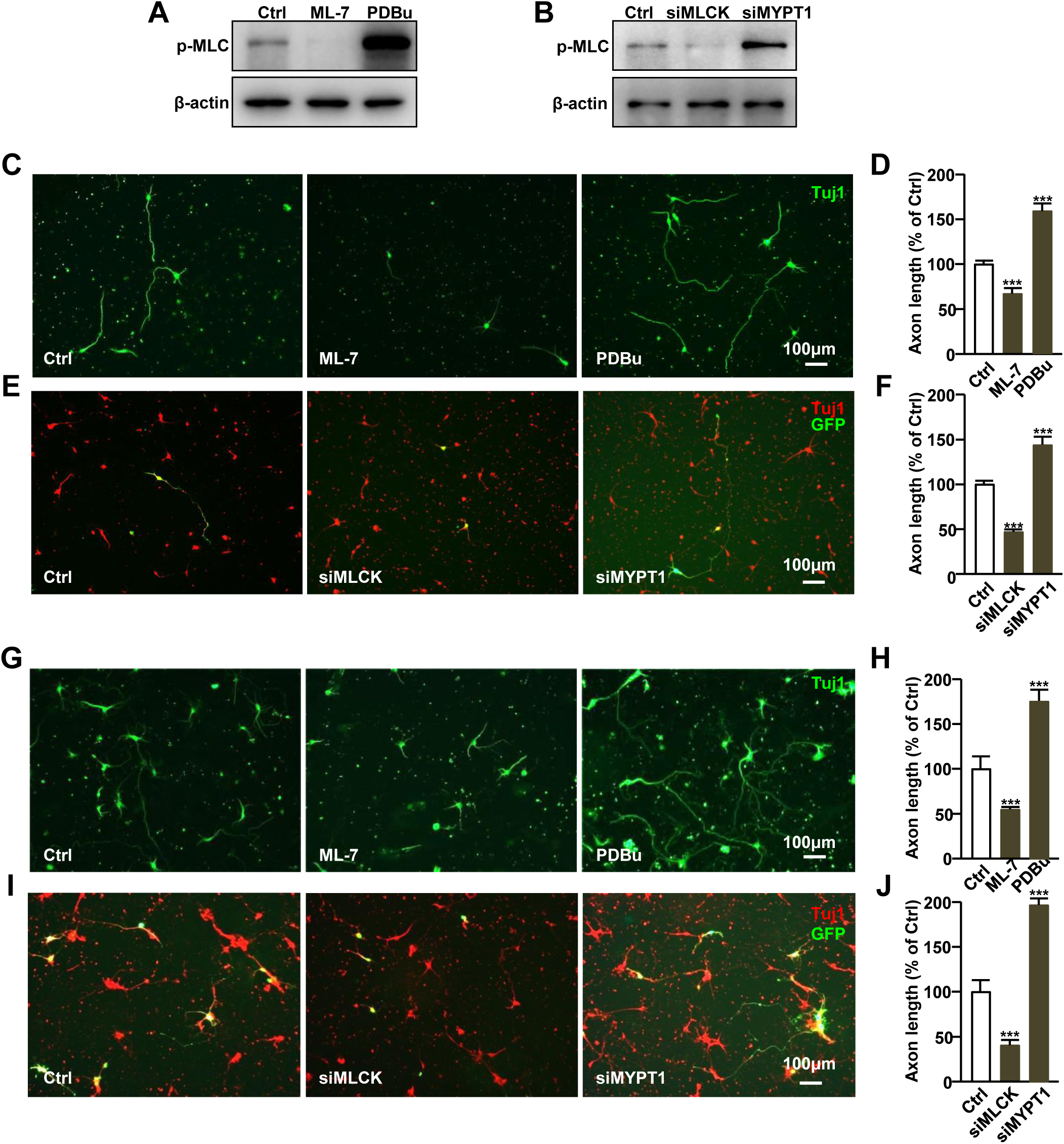
Inhibition of MLCP activity promotes axon growth from developing embryonic neurons. (A, B) Cortical neurons were isolated from embryonic day 14.5 embryos. Neurons were treated with DMSO, ML-7 or PDBu, or transfected with ctrl siRNA, siMLCK or siMYPT1. The p-MLC expression levels were examined using Western blot analysis after 3 days of culture. (C-F) coritcal neurons were treated with DMSO, ML-7 or PDBu or transfected with ctrl siRNA, siMLCK or siMYPT1. After three days, cells were stained with Tuj1 to visualize axons. Representative images (C, E) and quantification (D, F) of axonal length demonstrate ML-7 or siMLCK inhibits embryonic cortical neuronal axon growth. In contrast, inhibition of MLCP activity with PDBu or siMYPT1 promoted embryonic cortical neuronal axon growth (N=3, \*\*\**p* < 0.001). Scale bar: 100 μm. (G-J) Hippocampal neurons were isolated from embryonic day 18.5 embryos. Neurons were treated with DMSO, ML-7 or PDBu or transfected with scramble siRNA, siMLCK or siMYPT1. Similarly, Representative images (G, I) and quantification (H, J) of axonal length demonstrate that ML-7 or siMLCK inhibits hippocampal neuronal axon growth. In contrast, suppression of MLCP with PDBu or siMYPT1 promoted hippocampal neuronal axon growth (N=3, \*\*\**p* < 0.001). Scale bar: 100 μm.

### The local inhibition of MLCP activity at an injury site promoted optic nerve regeneration

Given the promising axon growth-promoting effects observed *in vitro*, we wondered whether inhibiting MLCP activity could stimulate *in vivo* CNS axon regeneration. The optic nerve forms part of the CNS and, like other neurons of the CNS, also have weak intrinsic axon-regenerating ability (Supplementary Figure S3). To test this, the right optic nerve of adult mice was crushed, and PBS or PDBu (50 μM) was applied locally to the lesion site using a gelatin sponge; alternatively, 2 μL of PDBu (50 μM) was microinjected into the vitreous body of another mice. Subsequently, 2 μL of Alexa Fluor 488-conjugated cholera toxin subunit B (CTB) was injected into the vitreous body to anterogradely label regenerating axons. After 3 days, the whole optic nerve was harvested, and CTB-labeled axon regeneration was assessed. The local administration of PDBu using a gelatin sponge resulted in numerous CTB-labeled axon fibers growing over the injury site, the longest being approximately 400 μm long (Figure 5A-C). In contrast, local PBS administration at the injury site or intravitreal PDBu injection did not significantly enhance axon regeneration beyond the injury site (Figure 5 A-C). This observation suggests that the only treatment employed in the injury site (the inhibition of MLCP activity within the growth cone) effective promote axonal growth.

**Figure 5.**
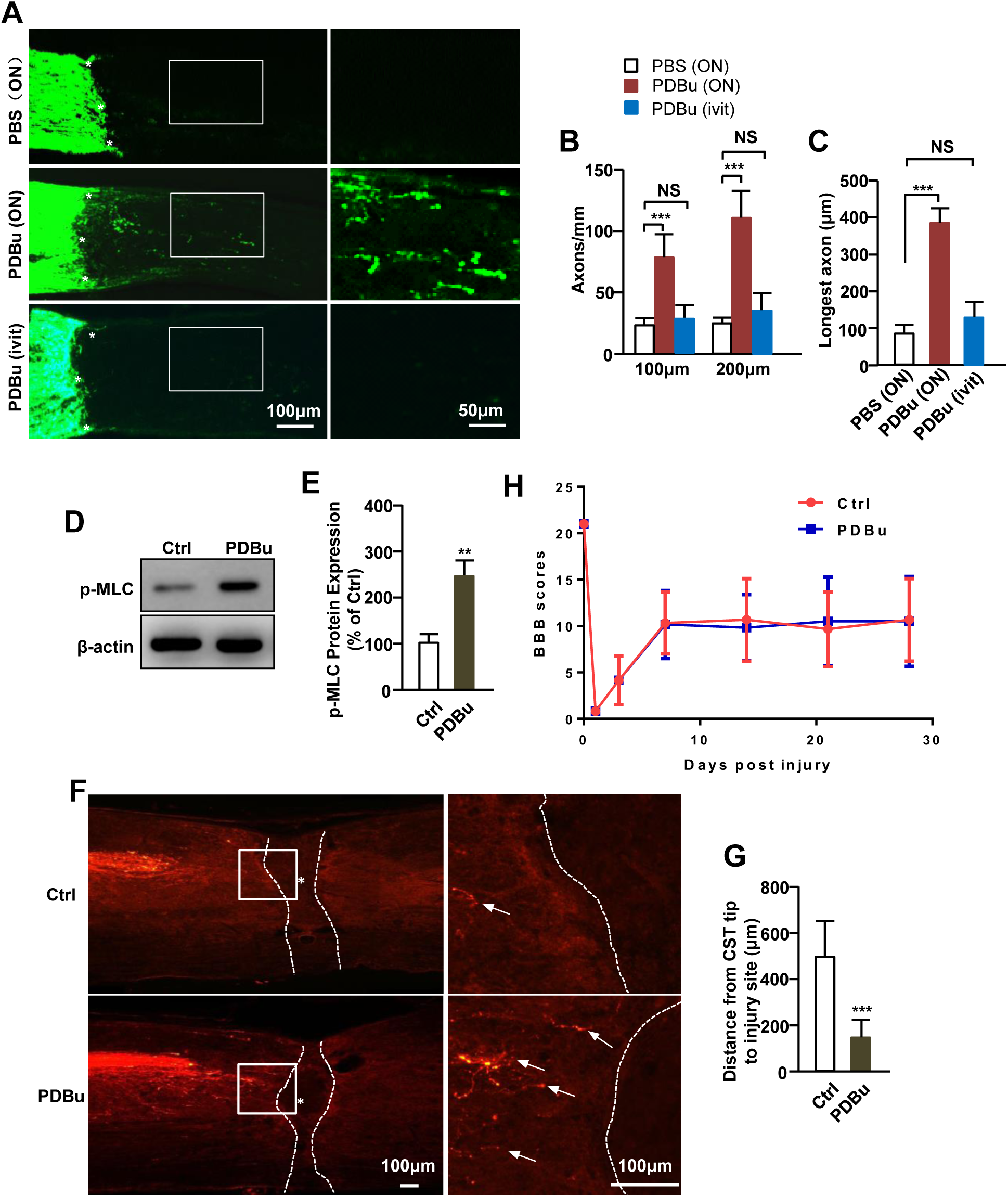
Inhibition of MLCP activity promotes CNS axon regeneration in vivo. (A-C) Local application of a gelatin sponge soaked with PBS (Ctrl) or 50.0 μM PDBu on the site of the lesion (ON), or microinjection of PDBu (2.0 μl, 50.0 μM) into the vitreous body (ivit) were performed after optic nerve crush injury. Three days later, CTB-labeled regenerating axons were quantified. Representative images of optic nerve sagittal sections (A) and quantification of CTB-positive fibers (B) at the point distance 100 μm and 200 μm from the injury site indicate that only local administration of PDBu at the site of the lesion promotes the axon growth. (C) The longest axons per section of group were measured and quantified. (D-H) T8 whole spinal cord crush injury was performed in adult mice then treated with PBS or 50.0 μM PDBu by intrathecal injection once every 4 days. BDA was injected into the right sensorimotor cortex 2 weeks after spinal cord injury to label the regenerating cortical spinal tract (CST) axons. After 2 weeks, sagittal sections of the spinal cord around the crush site were created, and the distance between the tip of the axons and injury site was measured. Representative image of Western blots (D) and its quantification (E) demonstrates that intrathecal injection of PDBu increases p-MLC level, it means that PDBu sufficiently inhibits MLCP activity. Representative image of spinal cord sagittal section (F) and the measurement of distance from BDA-positive axon tips to the injury site demonstrate that intrathecal injections of PDBu prevent injury-induced axon retraction. Arrow indicates BDA-positive axons, dash line indicates lesion site. (H) Mean BBB scores evaluation indicated that there was no difference between PDBu treatment and the PBS control group. (N=5, \*\*\**p* < 0.001).

### The local inhibition of MLCP activity prevented axonal retraction in the injured spinal cord

After spinal cord injury, corticospinal tract (CST) axons also did not regenerate spontaneously, principally because of the inhibitory environment (Supplementary Figure S4). To test whether local inhibition of MLCP activity can also promote spinal cord axon regeneration, mice were subjected to spinal cord crush injury at the level of the eighth thoracic vertebra (T8)^15^. Briefly, the T8 spinal cord was exposed and crushed for 1 second with jeweler’s forceps #5. A 100 μL PDBu (50 μM) or its vehicle, DMSO, was then administrated once every 4 days by intrathecal injection to block MLCP activity. After 2 weeks, 1.6 μL of 10 % biotinylated dextran amine (BDA) was microinjected into the sensorimotor cortex to label regenerating axons^16^. After a further 2 weeks, BDA-labeled regenerating axon was evaluated in the spinal cord section. Intrathecal PDBu injection significantly boosted p-MLC levels (Figure 5D, E) and prevented injury-induced axon retraction (Figure 5F, G), while also greatly reducing the number of retraction bulbs (Supplementary Figure S5). However, we did not observe any BDA-labeled axon fibers beyond the lesion site, and Basso–Beattie–Bresnahan (BBB) score evaluation following PDBu treatment did not differ from that of the control group (Figure 5H). Together, these results indicate that the local inhibition of MLCP activity reduces injury-induced retraction bulb formation and axon retraction in the spinal cord.

### MLCK may regulate axon regeneration independent of myosin II activity

We further examined whether the regulatory effect of MLCK/MLCP on axonal growth was mediated through myosin II activity. For this, dissociated DRG neurons were co-treated with blebbistatin (a pharmacological myosin II inhibitor) and ML-7. Consistent with our previous study ^1^, the inhibition of myosin II activity with 25 μM blebbistatin markedly promoted axonal growth (Figure 6A, B). Surprisingly, however, ML-7/blebbistatin co-treatment did not block the inhibitory effect of ML-7 on sensory axon regeneration (Figure 6A, B). Additionally, blebbistatin administration did not influence MLC phosphorylation levels in cultured DRG neurons (Figure 6C, D). Collectively, these results interestingly suggested that MLCK regulates axon regeneration may be independent of myosin II activity.

**Figure 6.**
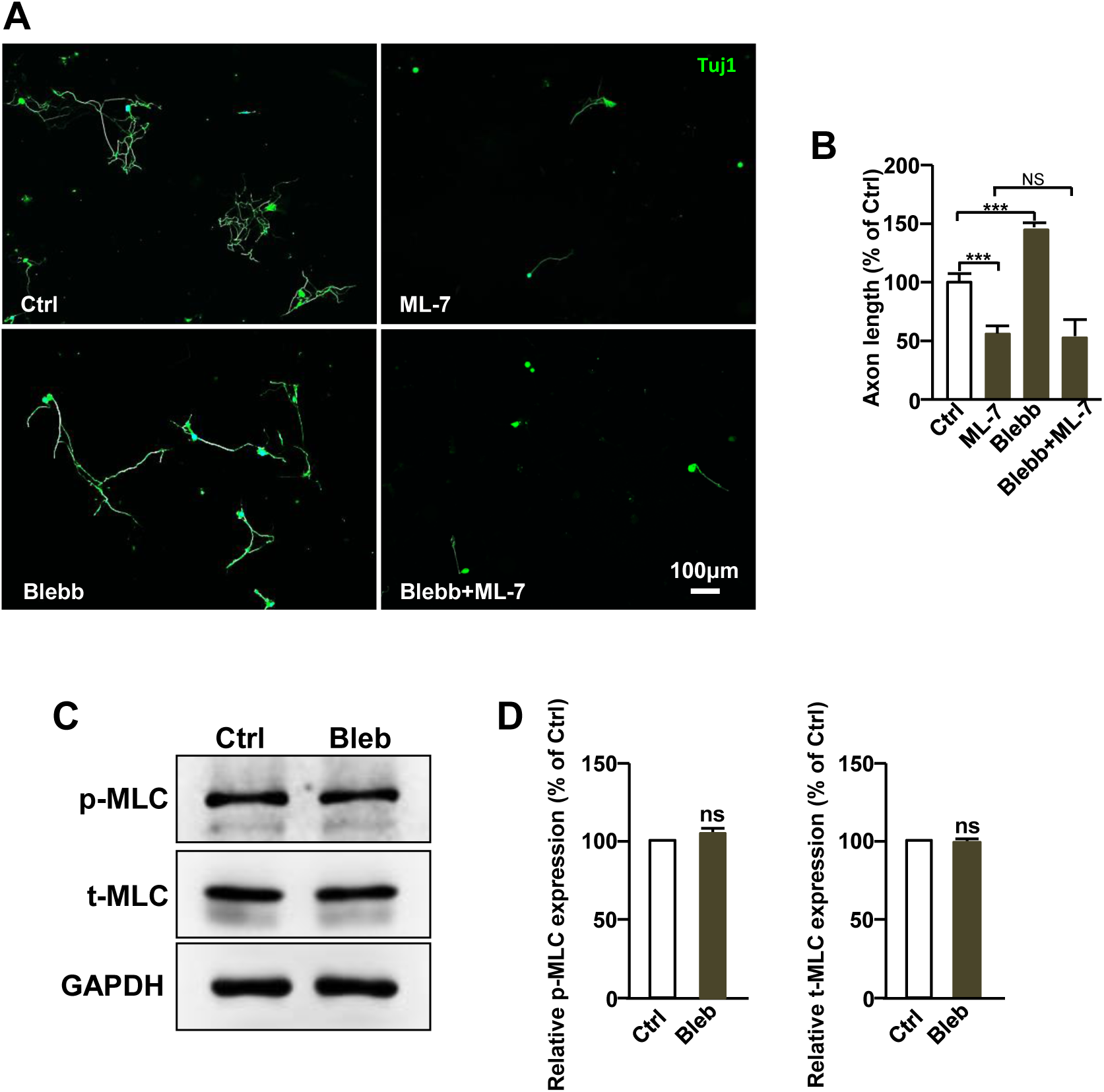
MLCK regulates axon regeneration independent of myosin II activity. (A, B) Adult L4-L5 DRG neurons were cultured *in vitro* for 3 days with the addition of the pharmacological inhibitors indicated. Representative images (A) and quantification of axon length (B) demonstrate that, although myosin II inhibitor blebbistatin markedly promoted sensory neuronal axon growth, however, it did not rescue the inhibitory effect of ML-7 on sensory axon growth (N=3, \*\*\**p* < 0.001). (C, D) The blebbistatin administration did not influence MLC phosphorylation levels in cultured DRG neurons, representative image of Western blots (C) and its quantification (D) (N=3, ns: no significant).

### Local F-actin distribution in the growth cone is regulated by MLCK and MLCP

Axonal regeneration requires both regulation of gene expression in the soma and the cytoskeletal assembly at the growth cone. We previously established a culture-and-replating model to investigate whether axonal regeneration was soma gene transcription-dependent or local growth cone cytoskeleton assembly-dependent^12,13^. Interestingly, only ML-7 treatment after replating was found to significantly inhibit axonal growth in peripheral sensory neurons. However, no effect was observed on axonal growth when ML-7 was administrated before cell replating (Figure 7A, B, C). Several studies have reported that peripheral axotomy triggers local cytoskeleton dynamics-dependent axon growth in DRG neurons ^13,16,20^. Consistent with the culture-and-replating results, axotomy-induced, local cytoskeleton assembly-dependent axon growth was also markedly inhibited by ML-7 treatment (Figure 7D, E). These results suggested that MLCK primarily influences local cytoskeleton dynamics during axonal growth. Accordingly, we further examined how MLCK/MLCP activity regulates growth cone cytoskeletal distribution during axonal growth. E14.5 cortical neurons were treated with ML-7 or PDBu for 3 days, following which microtubule and F-actin distribution in the growth cone was visualized using Tuj1 and phalloidin staining. The results indicated that ML-7 treatment increased growth cone size and F-actin content, whereas PDBu treatment exerted the opposite effect (Figure 8A-C). These findings may indicate that MLCK and MLCP activity regulates F-actin redistribution.

**Figure 7.**
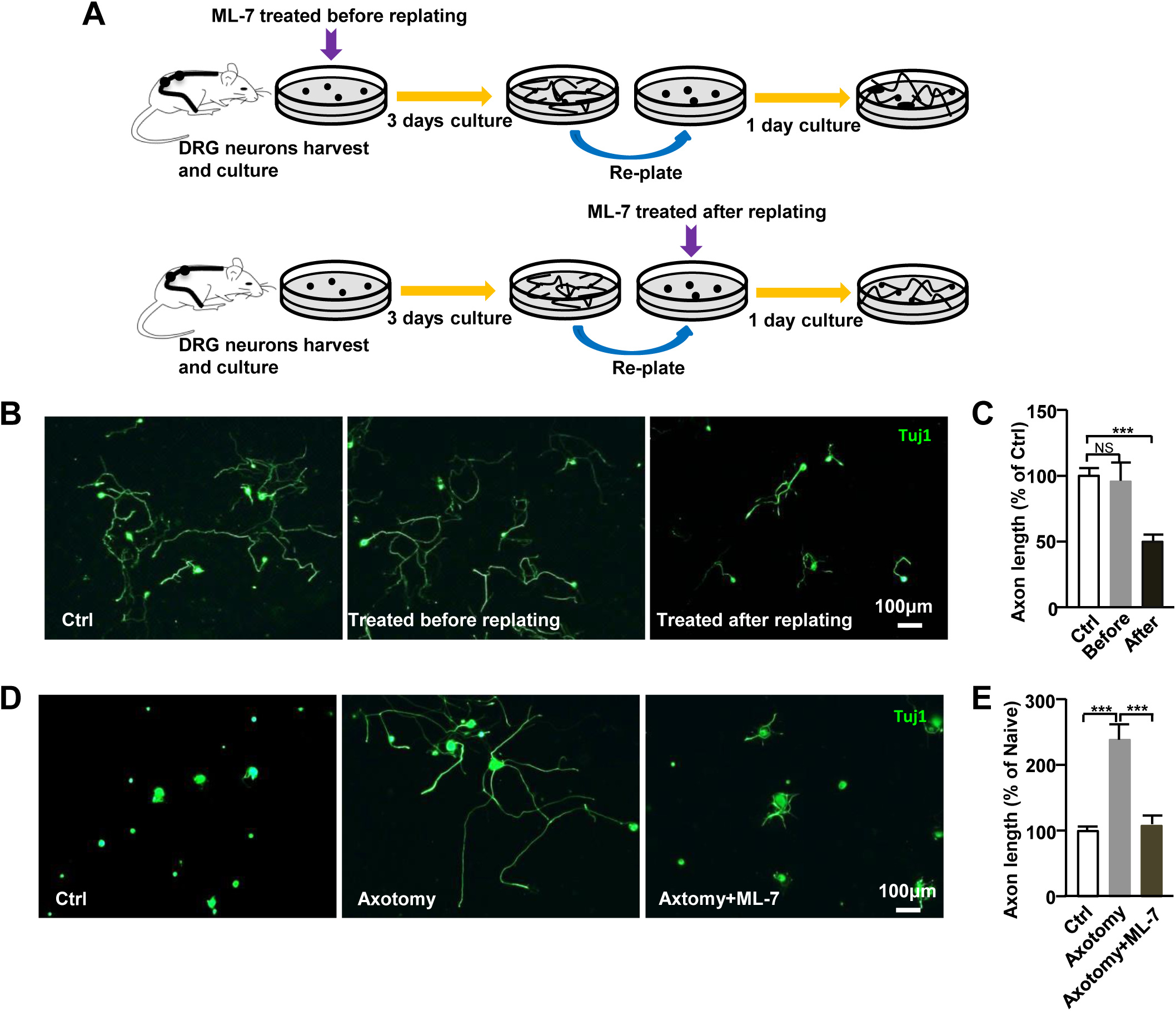
Inhibition of MLCK blocks transcription-independent axon growth of adult sensory neurons. (A) Schematic figure of the culture-and-replate model. Adult sensory neurons were cultured for 3 days then replated to allow growth of new axons for one day. During this process, ML-7 was added prior to or after replating. (B, C) Representative images (B) and quantification of axon length (C) indicate that treatment with ML-7 after replating significantly inhibits adult sensory axon growth. However, treatment with ML-7 prior to replating has no effect on adult sensory neural axon growth (N=3, \*\*\**p* < 0.001). (D, E) Mice underwent sciatic nerve axotomy 7 days previously. L4-L5 DRGs were harvested from naive or axotomized mice and cultured for 1 day with ML-7 treatment. Neurons were stained with Tuj1. Representative images (D) and quantification of axon length (E) indicate that blockade of MLCK activity with ML-7 inhibits axotomy-induced axon growth of adult sensory neurons (N=3, \*\*\**p* < 0.001).

**Figure 8.**
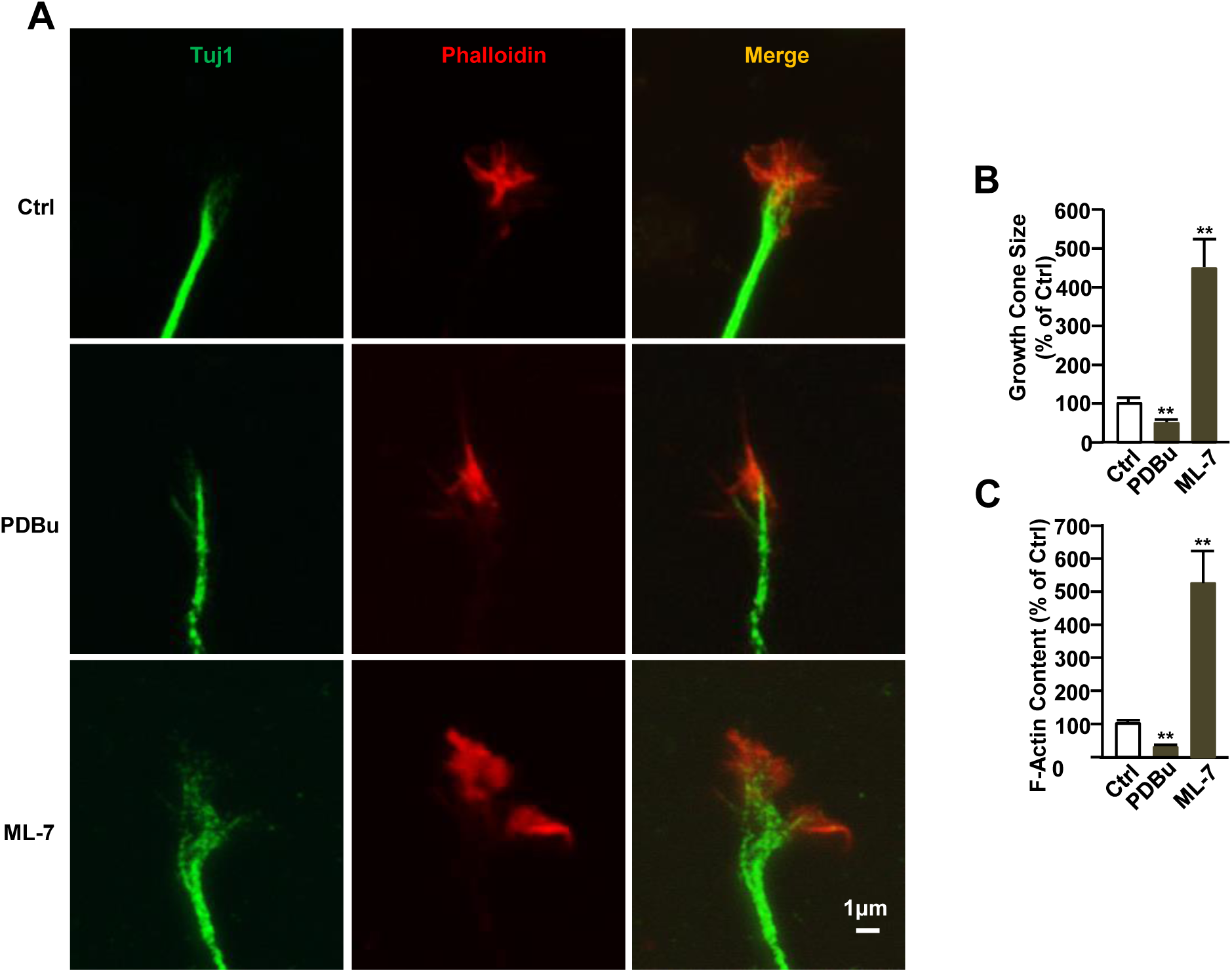
MLCK and MLCP activity regulate F-actin distribution of the growth cone. (A, B) Embryonic E14.5 cortical neurons were isolated and treated with DMSO, PDBu or ML-7. After 3 days of culture, neurons were stained with Tuj1 antibody (Green) to visualize tubulin and phalloidin (Red) to visualize F-actin. Representative images (A), quantification of the size of the growth cone (B), and F-actin content (C) indicate that ML-7 treatment increases the growth cone size and F-actin content, and PDBu treatment reduces the growth cone size and the F-actin content. Scale bar: 1μm (N=3, \*\*\**p* < 0.001).

## Discussion

In the present study, we demonstrated that peripheral axotomy increases MLCK and MLC phosphorylation levels in adult DRG neurons subsequently. Thus, when MLCK activity was suppressed in mature DRG neurons *via* ML-7- or specific siRNA, axonal growth was blocked in treated neurons both *in vitro* and *in vivo*. Conversely, an inhibit in MLCP activity *via* the knockdown of MYPT1 significantly promoted axonal growth in both PNS and CNS neurons. We further found that the MLCK and MLCP primarily regulate F-actin redistribution in the growth cone during the axon regeneration process.

Axon injury leads to disconnection between neurons and their targets, thereby destroying functional circuits and behavioral outputs. This implies that for the successful restoration of impaired nervous system function, injured axons must regenerate to their original targets and reconstruct functional circuits. Interestingly, it has been reported that the coordinated regulation of the neuronal cytoskeleton dynamics is essential for axon shaft extension ^21^. Among them, the microtubule and actin are two major components of neuronal cytoskeleton that involves axonal extension. For example, Athamneh et al. found that the speed of axon elongation is highly correlated with the bulk forward translocation rate of microtubules ^22^. In addition, it also has been reported that the axon extension depends on actin assembly and microtubule-actin interactions^23^. Unsurprisingly given these observations, increasing attention has been paid to promoting axon regeneration *via* the modulation of local cytoskeletal dynamics in the growth cone. Myosin II is a key cytoskeleton protein, which activated through the MLCK-mediated phosphorylation of its light chain and deactivated *via* the dephosphorylation of the N-terminus of its light chain by MLCP. Here, we found that the MLCK expression and phosphorylation level of MLC was increased in adult DRG neurons after sciatic nerve injury, while the inhibition of MLCK blocked axon regeneration. These results indicate that activated MLCK is required for mammalian axon regeneration. If thus, an increase of MLCK expression will activate myosin II during axon regeneration. However, in our previous study, we found that pharmacological blockade of myosin II activity with blebbistatin stimulated axon growth ^1^, which suggested that myosin II plays an inhibitory role in this process. This apparent contradiction might be attributed to blebbistatin being a specific inhibitor of non-muscle myosin II ATPase activity and may not affect MLCK activity or the phosphorylation status of MLC^24^. Indeed, blebbistatin treatment did not influence MLC phosphorylation levels in our study, in line with a previous report in which it was found that blebbistatin does not alter the phosphorylation status of MLC in sensory neurons and trabecular meshwork cells^24,35^. The reported mechanism of blebbistatin is not through competition with the ATP binding site of myosin. Instead, it selectively binds to the ATPase intermediate state associated with ADP and inorganic phosphate, which decelerates the phosphate release. Importantly, blebbistatin does not impede myosin’s interaction with actin or the ATP-triggered disassociation of actomyosin^36^. It rather inhibits the myosin head when it forms a product complex with a reduced affinity for actin. This indicates that blebbistatin functions by stabilizing a particular myosin intermediate state that is independent of the phosphorylation status of MLC. Additionally, our results demonstrated that blebbistatin treatment could not rescue the inhibitory effect of ML-7 on axon regeneration. If MLCK regulates axon growth through the activation of myosin, the inhibitory effect of ML-7 on axon growth might be influenced by blebbistatin, a NMII inhibitor. However, our findings reveal that the combination of blebbistatin and ML-7 does not alter the rate of axon outgrowth compared to ML-7 alone. This suggests that the roles of ML-7 and blebbistatin in axon growth are independent. It means MLCK may regulate axon growth independent of NMII activity. Interestingly, it has been reported that MLCK regulates cell migration not by MLC phosphorylation, but possible MLCK may serve as an F-actin-binding protein stabilizing the F-actin/myosin II network of the membrane cytoskeleton^25^. Additionally, recent study also identified multiple noncanonical targets of MLCK through CRISPR-Cas9/phosphoproteomics method^26^, thus it is also possible that MLCK regulates axon growth its noncanonical targets or unknown mechanism. They also found that MLCK plays an important role in vasopressin-induced actin depolymerization^26^. Studies have shown that the combination of MLCK with F-actin is critical to its function. Short-chain MLCK binds to F-actin via three DFRXXL motifs at its N-terminal, while long-chain MLCK has two additional DFRXXL motifs and six IG-like modules, which may give it unique binding properties in cell localization^25^. During axon growth, the dynamic changes of F-actin are regulated by a variety of proteins, such as ADF/cofilin, Arp2/3, Eps8, Profilin, myosin II, myosin V, etc. These proteins regulate axon growth and orientation by influencing the polymerization and depolymerization of F-actin. For example, cofilin increases the availability of actin by promoting the depolymerization of F-actin, which promotes axon growth^37^. Furthermore, MLCP inhibition also increase actin arc movement in growth cone^5^. This means that MLCK/MLCP may also regulate axon growth via actin dynamics. In addition, other studies also have shown that the absence of MLCK not only impairs the synthesis of transmembrane complexes, such as integrin, and fibronectin, but also disrupts the balance of membrane tension and protrusion, thereby causing the membrane cytoskeleton instability^38, 39^. We think these may affect axon growth.

Meanwhile, adult sensory neurons express different isoforms of myosin II, which also exert distinct effects on axonal regeneration^27^. For example, the knockdown of myosin IIA enhances axonal growth, while the knockdown of myosin IIB increases F-action retrograde flow and inhibits axon growth^1,28–31^. A study in Neuro-2A cells also revealed that myosin IIA is involved in axon retraction, whereas myosin IIB was reported to contribute to axon extension^28,29^. Interesting, in neurons, although expression of both NMIIA and NMIIB has been reported, the NMIIB being the predominant one^40^. Under physiological conditions, MLCP is activated by PKC, and it has been reported that the PKC-mediated activation of myosin II is restricted to the T zone of growth cones; the results of the present study suggest that growth cones require low-to-moderate levels of myosin II activity and F-actin for best growth rates. This observation emphasizes the complexity of the role of myosin II in axonal growth, the precise molecular mechanisms of which require further in-depth investigation.

The inability of neurons in the CNS to regenerate their axons after injury is attributable, at least in part, to the presence of inhibitory molecules at the injury site, including CSPGs and myelin. Here, we found that the inhibition of MLCP activity with PDBu or the siRNA-mediated knockdown of MYPT1 promoted axonal growth in adult sensory neurons over both CSPG and myelin substrates. Our findings also further indicated that the local inhibition of MLCP by PDBu not only induces axon regeneration in peripheral sensory neurons but also promotes axon regeneration in the optic nerve. Studies have shown that MLCK possesses several actin-binding domains to which F-actin can bind and form bundles^32,33^ and that directional growth cone motility during axon growth is controlled by actin-based machinery. Growth cone movement during axon growth is achieved through protrusion towards attractive guidance cues and retraction from repulsive ones in an actin-dependent manner. In the classic molecular clutch model, strong mechanical coupling between the F-actin cytoskeleton and the substrate, mediated by cell adhesion complexes, transmits the forces generated by the cytoskeleton into rearward traction forces, enabling the forward movement of the growth cone. Our results indicated that treating DRG neurons with ML-7 induces the increases F-actin content in the growth cone. Conversely, the inhibition of MLCP following PDBu administration resulted in decreased F-actin content. Our findings are consistent with the results of a previous study on Helisoma, in which treatment with ML-7 increased F-actin content at the C/P-domain interface, thus inhibiting axonal regeneration^34^. This suggests that MLCK or MLCP may control mammalian axonal regeneration by regulating growth cone F-actin distribution. Together, our findings indicate that directly targeting growth cone cytoskeleton components may be a promising strategy for promoting mammalian axon regeneration.

## Statements & Declarations

### Author Contribution

Saijilafu, W.H. W and J.J. M designed the experiment. W.H. W and J.J. M performed the experiments and analyzed the data. W.H. W, Y.Y, Y.X. M and Saijilafu co-wrote the paper with all authors’ input.

### Ethics approval

The animal experiments were conducted in accordance with the National Institutes of Health Guide for the Care and Use of Laboratory Animals (NIH Publications No. 8023, revised 1978) and were approved by the Ethics Committee of the First Affiliated Hospital of Soochow University.

### Consent to publish

All the authors consent for publication.

### Data availability

All data generated or analyzed during this study are available from the corresponding author upon reasonable request.

---

**Figure S1.**
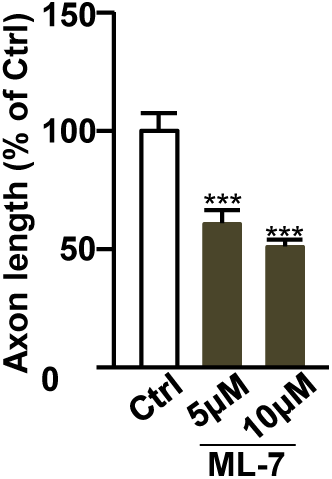
ML-7 treatment inhibits axon growth in a dose-dependent manner. The DRG neurons from adult mice were cultured and treated with a series of doses of ML-7. After 3 days culture, neurons were stained with Tuj1 antibody and axonal lengths quantified.

**Figure S2.**
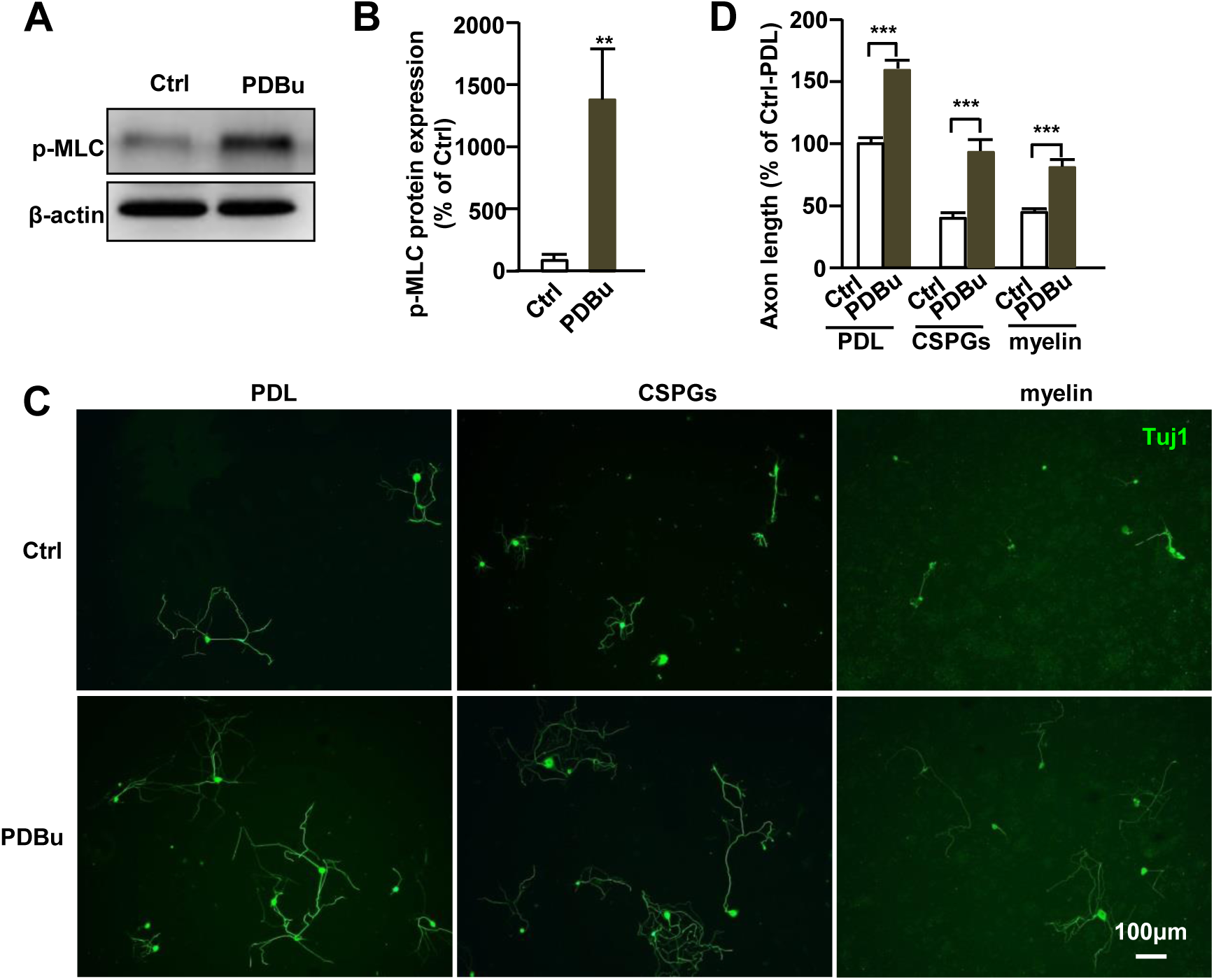
PDBu treatment promotes axon growth from adult peripheral sensory neurons. (A, B) Representative Western blot image (A) and quantification of protein band (B) indicates that administration of PDBu, the MLCP pharmacological inhibitor, in cultured DRG neurons increases p-MLC expression (N=3, ** *p* < 0.01). (C, D) DRG neurons were cultured on permissive (PDL) or inhibitory substrates (CSPGs or myelin) in the presence or absence of PDBu, as indicated. Neurons were stained with Tuj1. Representative images (C) and quantification of axon length (D) show that increased p-MLC expression with PDBu addition, which promotes adult sensory neuronal axon growth on either permissive substrate PDL or inhibitory substrates, such as CSPGs and myelin. (N=3, \*\*\**p* < 0.001).

**Figure S3.**
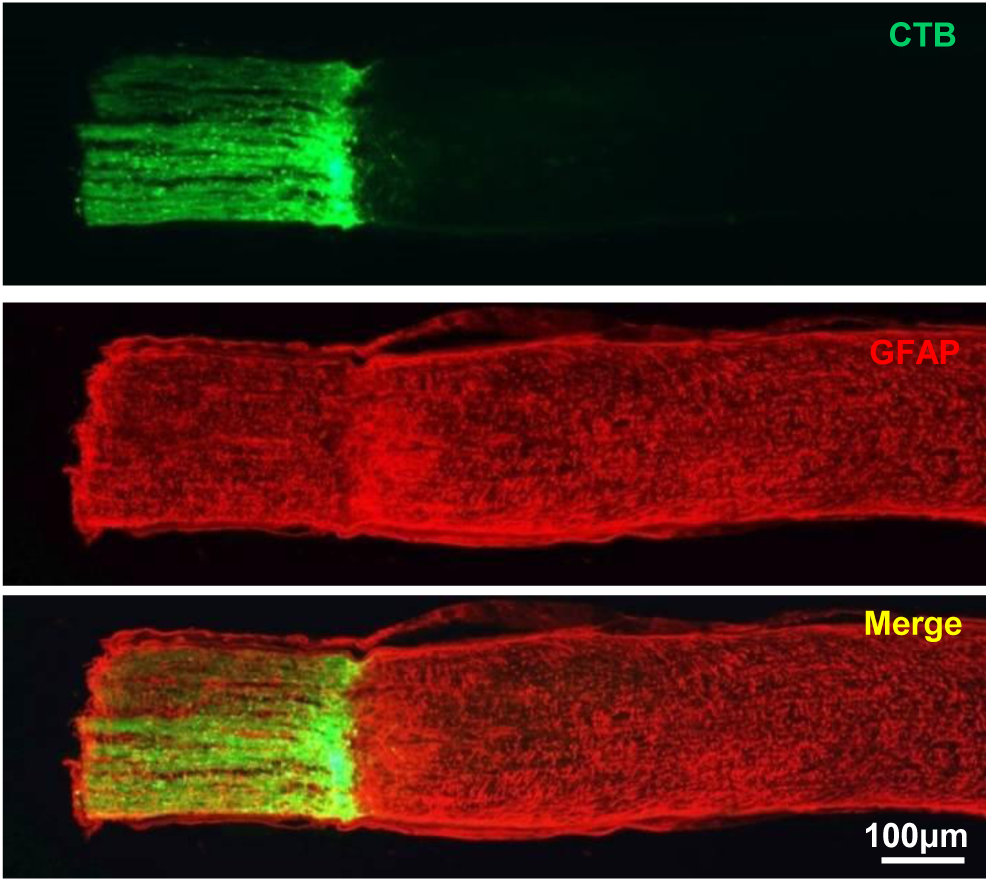
Optic nerve crush injury model. Optic nerve was crushed behind the eyeball with a custom-modified jeweler’s 5^#^ forceps for 1 second. Two weeks later, 2.0 µl Alexa Fluor 488-conjugated CTB (green) was injected intravitreally to label regenerating optic nerve axons. Immunofluorescent staining of GFAP staining (red) was also performed in the longitudinal section of crushed optic nerves.

**Figure S4.**
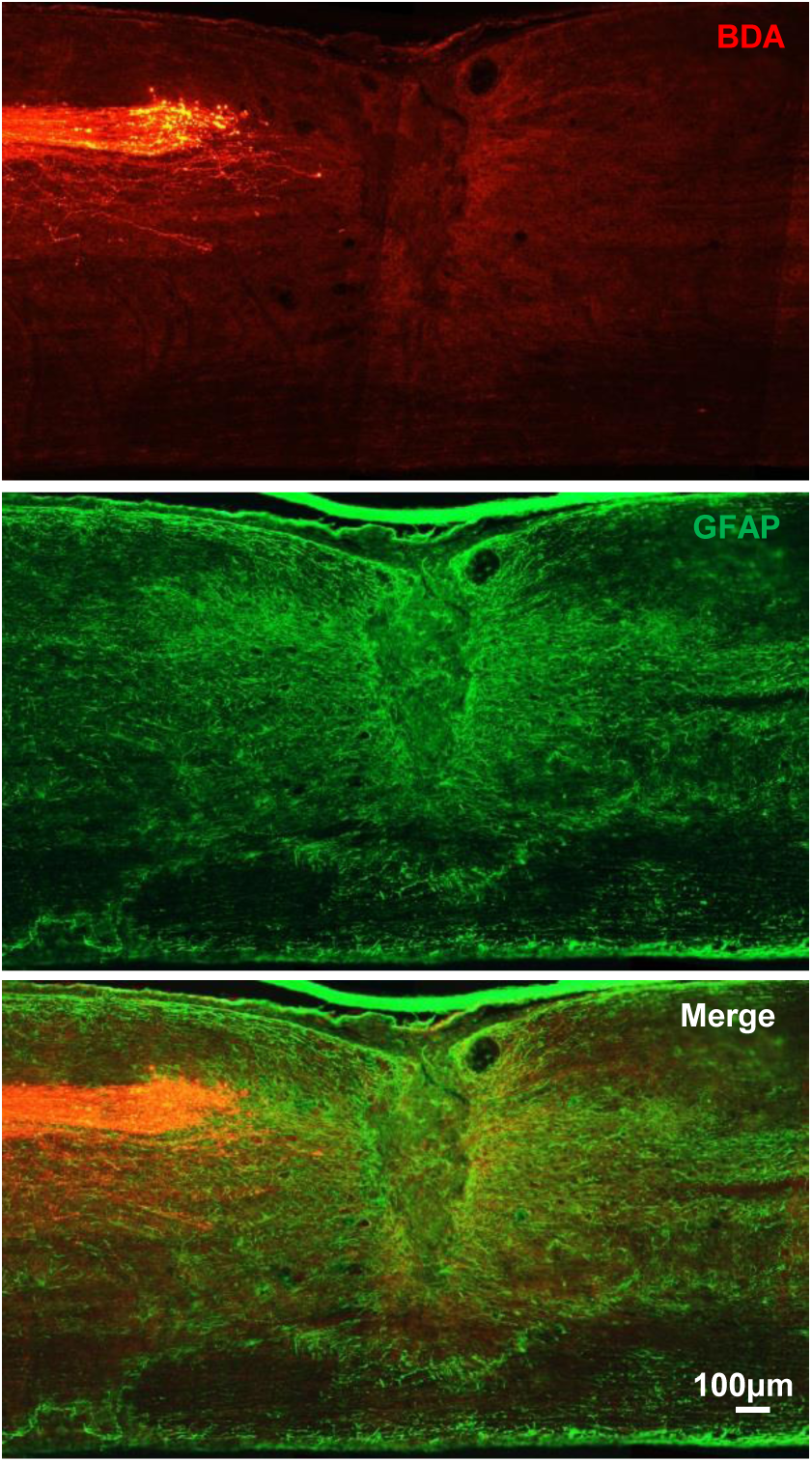
Whole spinal cord crush injury model. The whole spinal cord was crushed at T8 level with a custom-modified jeweler’s 5^#^ forceps for 1 second. Two weeks later, a 1.6 μl aliquot of 10% BDA was injected into the sensorimotor cortex to label regenerating cortical spinal tract axons, and another two weeks later, the spinal cord was dissected out. Immunofluorescent staining of BDA (red) and GFAP staining (green) was performed in the sagittal section of spinal cord.

**Figure S5.**
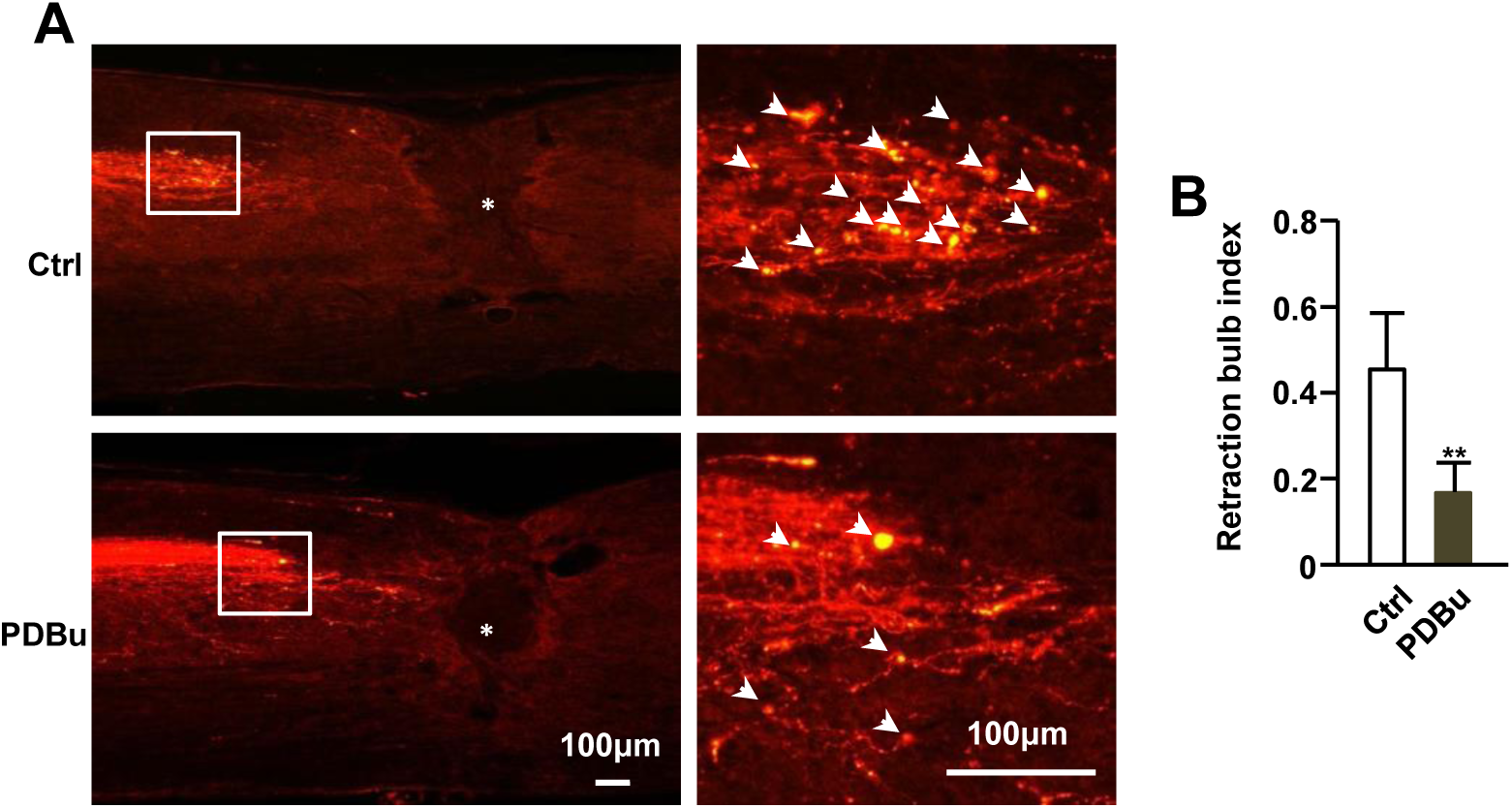
Intrathecal PDBu injection prevented axonal retraction bulb formation. Representative images of spinal cord sagittal section (A) and quantification of retraction bulb formation (B). White triangle indicates retraction bulb (N=3, *** *p* < 0.001).

